# COCOMO2: A coarse-grained model for interacting folded and disordered proteins

**DOI:** 10.1101/2024.10.29.620916

**Authors:** Alexander Jussupow, Divya Bartley, Lisa J. Lapidus, Michael Feig

## Abstract

Biomolecular interactions are essential in many biological processes, including complex formation and phase separation processes. Coarse-grained computational models are especially valuable for studying such processes via simulation. Here, we present COCOMO2, an updated residue-based coarse-grained model that extends its applicability from intrinsically disordered peptides to folded proteins. This is accomplished with the introduction of a surface exposure scaling factor, which adjusts interaction strengths based on solvent accessibility, to enable the more realistic modeling of interactions involving folded domains without additional computational costs. COCOMO2 was parameterized directly with solubility and phase separation data to improve its performance on predicting concentration-dependent phase separation for a broader range of biomolecular systems compared to the original version. COCOMO2 enables new applications including the study of condensates that involve IDPs together with folded domains and the study of complex assembly processes. COCOMO2 also provides an expanded foundation for the development of multi-scale approaches for modeling biomolecular interactions that span from residue-level to atomistic resolution.

**Table of Contents Figure:** 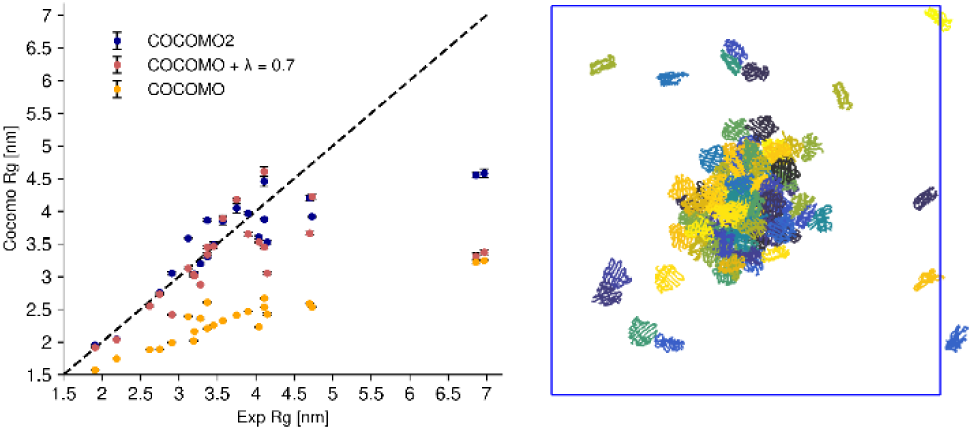

## INTRODUCTION

Intermolecular interactions between biomolecules play a central role for many aspects of biology. Specific interactions lead to oligomerization^1^, aggregation^2^, and the formation of large assemblies such as the ribosome^3^, the mediator complex^4^, the nuclear pore complex^5^, virus capsids^6^, or bacterial microcompartments^7^. In the crowded cellular interior, non-specific interactions are inevitable, leading to clustering^8^ or phase separation^9^. Biomolecular phase separation, particularly liquid-liquid phase separation (LLPS), plays a crucial role in the formation of membrane-less organelles within cells, driving the compartmentalization of essential biological processes. These condensates, which include organelles such as stress granules^10^, nucleoli^11^, and P-bodies^12^, form through a dynamic and reversible process involving proteins, RNA, and other biomolecules^13,14^. Understanding the molecular mechanisms behind complex assembly, dynamic clustering, and phase separation is key to uncovering how cells organize biochemical function in space and time^13,15^.

To complement experiments, computational tools like molecular dynamics (MD) simulations, both atomistic^16,17^ and coarse-grained^18–20^, have been employed to study biomolecular interactions and the formation of higher-order structures. Atomistic simulations provide detailed representations of every atom in a biomolecule, offering precise insights into molecular interactions. However, their high computational cost limits their applicability to small systems and short-time scales (typically not exceeding microsecond scales)^21^. Coarse-grained models overcome these limitations by simplifying the complexity of biomolecules, thus enabling simulations of larger systems over longer time scales^22,23^ and making them particularly suited for studying concentration-dependent phase separation, condensate formation^18,19^ and biomolecular complex assembly processes^24,25^.

The original COCOMO (Concentration-dependent Condensation Model)^26^ was developed as a one-bead-per-residue coarse-grained model specifically capturing key interactions between intrinsically disordered proteins (IDPs) and RNA that drive phase behavior. However, folded proteins and multi-domain proteins (MDPs) often also play an active role in liquid-liquid phase separation^27–29^ and are the main components of complex assemblies. This leads to a need for a revised model that is suitable for both IDPs and proteins with folded domains.

Here, we present COCOMO2, an improved version of the original COCOMO force field that extends its applicability to folded and multi-domain proteins. Building on the original COCOMO model^26^, we modeled folded domains via elastic network restraints and introduced a surface exposure scaling factor λ to modulate the interaction strength of residues based on their degree of solvent accessibility. This adjustment ensures that buried and partially exposed residues contribute less to intermolecular interactions than surface-exposed residues, effectively capturing solvation effects but without the additional expense of an implicit solvent model^30,31^. We also refined the approach for determining saturation concentrations (*c_sat_)*^32^ from simulations and introduced a computationally efficient protocol for approximating *c_sat_* via the potential energy of condensate structures based on theory^33–35^.

This allowed us to parameterize COCOMO2 directly using *c_sat_* data obtained either from LLPS or solubility experiments, unlike from other models that rely primarily on matching single-chain properties^19,20,36–38^, including the recent expansion of CALVADOS for multi-domain proteins^36^. In contrast to CALVADOS3, which assigns individual parameters to each amino acid, COCOMO2 provides a simpler model with fewer parameters by grouping polar and hydrophobic residues. COCOMO2 demonstrates significant improvements in accuracy for both phase separation behavior and single-molecule properties compared to the original COCOMO model, providing a more versatile framework for studying phase separation and solubility phenomena in diverse cellular contexts, including IDPs, MDPs, and RNA(-protein) condensates. While primarily focused on non-specific interactions, COCOMO2 could also be applied to model specific interactions involved in complex assemblies by adding system-specific interaction terms.

## METHODS

### Coarse-Grained Model

COCOMO2 builds on the structure of the original COCOMO model. Each amino acid or RNA nucleotide residue is represented as a single spherical particle. The total interaction energy in the system is defined as:

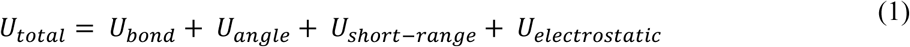

*U*_*bond*_ represents the bonded potential, with a harmonic bond potential ensuring connectivity along the chain:

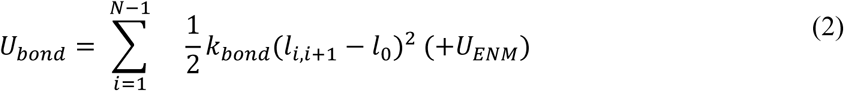

where *l*_*i*,*i*+1_is the distance between two neighboring residues, *k_bond_* =·4184 kJ/(mol·nm^2^) is the bond constant, and *l_0_* is the equilibrium bond length. *l_0_* is set to 0.38 nm for proteins, the average C_α_–C_α_ distance, and to 0.5 nm for nucleotides, the average backbone distance for single-stranded nucleic acids^39^.

For folded domains, an additional elastic network model (ENM) is applied to stabilize the higher-order structural elements:

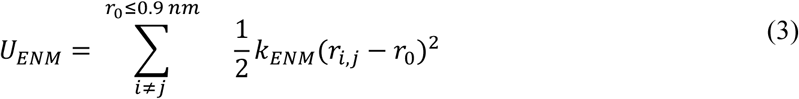

with *r*_*i*,*j*_ as the distance between two beads in a folded domain, *k_ENM_* =·500 kJ/(mol·nm^2^) as the force constant of the ENM, and *r*_0_as the equilibrium distance based on the initial reference conformation. Only residue pairs with a *r*_0_ ≤ 0.9 *nm* are considered for the ENM. Additionally, consecutive residues are excluded as they are already accounted for with the bonded potential.

The angle potential *U*_*angle*_ maintains the chain stiffness:

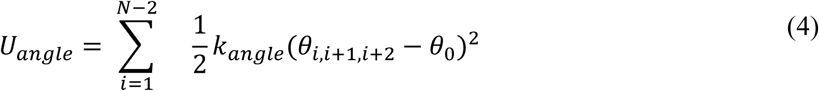

with λ_*i*,*i*+1,*i*+2_ as the angle between three consecutive beads, *k*_angle_ =·4.184 kJ/(mol·rad^2^) as the angle constant for proteins and 4.184 kJ/(mol·rad^2^) for nucleic acids, and λ_0_= 180° as the target angle.

The pairwise nonbonded short-range 10–5 Lennard-Jones potential (*U*_short-range_) is slightly modified in COCOMO2 compared to the original COCOMO:

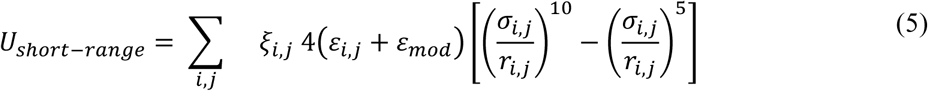

where *r*_*i*,*j*_ is the inter-particle distance. σ_*i*,*j*_ = 0.5(σ_*i*_ + σ_*j*_) is the distance at which the potential is zero, The effective radii σ_*i*_ were set as σ_*i*_ = 2*r*_*i*_ · 2^−1/6^, where *r*_*i*_ is the radius of a sphere with equivalent volume of a given residue. 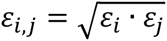 is the depth of the potential well. ε_mod_ is added to either enhance interaction between positively charged residues (Arg, Lys) and aromatic residues (Phe, Tyr, Trp) (ε_*R*/*K*−*F*/*Y*/*W*_ = 0.3 kJ/mol), between positively charged residues and nucleotides (ε_*R*/*K*−nucleic_ = 0.2 kJ/mol) or, new in COCOMO2, between aromatic residues (ε_*F*/*Y*/*W*−*F*/*Y*/*W*_ = 0.1 kJ/mol). 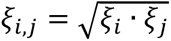 is the interaction strength. For disordered regions, the interaction strength (ξ) remains constant at 1, while for residues in the folded domains, ξ is scaled based on **Eq. 6**.

The surface exposure scaling factor λ directly influences the interaction strength ξ, modulating electrostatic interactions based on solvent accessibility:

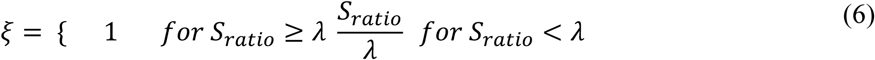

with *S_ratio_* = *S*_residue_/*S_ref_* as the ratio between the surface area of a residue (**S*_residue_*) and an amino-acid-specific reference area (*S*_ref_). *S*_ref_ was calculated as the surface area of an amino acid embedded in alanine α-helix and is reported in **Table S1** (in nm^2^). The interaction strength is always 1 for residues in disordered regions, while for residues in folded domains, it decreases as a function of *S*_ratio_, reaching zero for fully buried residues. 𝜉 is calculated based on the initial reference structure and is not updated during the simulation.

Electrostatic effects are described with an adjusted Debye–Hückel potential (𝑈*_electrostatic_*):

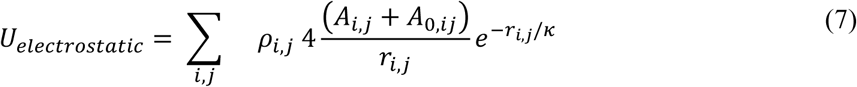

where *r_i,j_* is the inter-particle distance. *A*_i,j_ = *A*_i_ · *AA*_j_ reflect the attractive or repulsive electrostatic interactions, with 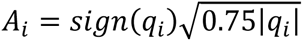 calculated from residue charges q (see ^40^). *A*_0,i,j_ = *A*_0,i_ · *A*_0,j_ reflects the effective repulsion due to solvation effects. The Debye screening length κ is set to 1 nm, corresponding to an ionic strength of ∼100 mM.

For COCOMO2, the following parameters were optimized: ε*_polar_*, ε*_hydrophobic_*, *A*_0,polar_, *A*_0,*hydrophobic*_, λ, with Arg, Asn, Asp, Cys, Gln, Glu, His, Lys, Ser, and Thr as polar residues, and Ala, Gly, Ile, Leu, Met, Phe, Pro, Trp, Tyr, and Val as hydrophobic residues. **Table 1** shows a comparison between the original COCOMO and the new COCOMO2 parameters.

**Table 1:**
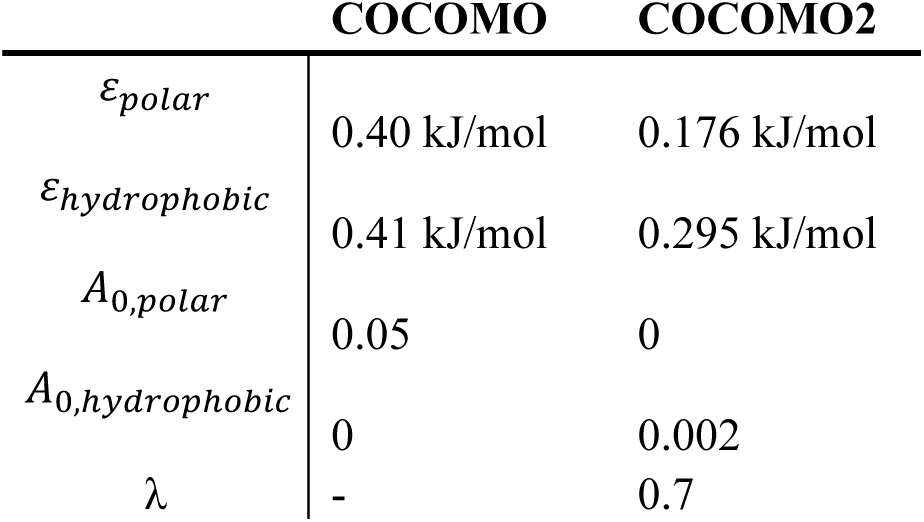
Summary of the COCOMO and COCOMO2 parameters.

|  | COCOMO | COCOMO2 |
| --- | --- | --- |
| $\epsilon_{polar}$ | 0.40 kJ/mol | 0.176 kJ/mol |
| $\epsilon_{hydrophobic}$ | 0.41 kJ/mol | 0.295 kJ/mol |
| $A_{0,polar}$ | 0.05 | 0 |
| $A_{0,hydrophobic}$ | 0 | 0.002 |
| $\lambda$ | - | 0.7 |

### Molecular Dynamics simulations

All molecular dynamics simulations with COCOMO2 were performed using OpenMM 8.0.0^41^. Langevin dynamics was applied with a friction coefficient of 0.01 ps^-1^, and simulations were run as an NVT ensemble at 298 K. The integration time step was set to 10 fs during equilibration and to 20 fs for production runs. Simulations were conducted under periodic boundary conditions with nonbonded interactions truncated at 3 nm. Residues separated by one bond were excluded from nonbonded interactions energy calculation. With these parameters, it takes 4 hours to sample 1 µs for a system of 180 randomly distributed 166-residue IDPs (29880 beads in total) in a 100 nm box on an RTX 2080 Ti GPU card. For a fully condensed system, the time increases to 11 hours due to an increased number of nonbonded interactions below 3 nm.

Single-chain simulations were performed for 23 multidomain proteins, using the same starting structures as those used in the CALVADOS expansion for multi-domain proteins^36^. Systems were equilibrated for 5000 steps and followed by 1 µs production runs. Details of the systems, simulation box sizes and folded domain region definitions are provided in **Table S2**. The radii of gyration (*R_g_*) were calculated using the MDtraj library^42^ and the *SS*_residue_ values were determined using the Gromacs SASA tool^43,44^. Multi-chain simulations typically involved 100 to 300 chains, with box sizes between 50 and 200 nm. Production runs were generally conducted for 5 μs each, with detailed simulation conditions, including folded domain definitions, listed in **Tables S3-S7**. Protein systems with folded domains were taken from Golovanov *et al.*^45^ or Cao *et al*.^36^. For estimating saturation concentration (*c_sat_*), simulations were set up as a mixture of pre-formed condensate and monomers to ensure smoother convergence and to avoid hysteresis effects (see below). All visualizations were done with VMD^46^.

### Saturation concentration estimation

In COCOMO, the *c_sat_* values were estimated as the highest concentration at which condensate formation was not observed during the simulation time, while the critical concentration (*c_crit_)* was defined as the lowest concentration at which condensate formation was observed. Alternatively, one can determine *c_crit_* by starting simulations from a pre-formed condensate to find the minimum concentration at which a condensate remains stable.

In the absence of supersaturation effects, *c_sat_* should approach *c_crit_* given a sufficiently long timescale, and *c_sat_* is expected to be the concentration of the dilute phase when there is coexistence with condensates reached either from the disperse phase or a pre-formed condensate. However, determining c_sat_ or c_crit_ via simulations started from the disperse phase may lead to overestimated saturation concentrations, whereas simulations started from a condensate may underestimate the saturation concentration (**Fig. 1**). This is due to the time it takes to nucleate condensation or melting a condensate. Since nucleation is a stochastic process, nucleation events may not occur within the simulation timescale^14^, particularly at concentrations just above *c_sat_* or at very low concentrations when interactions do not occur frequently.

**Figure 1.**
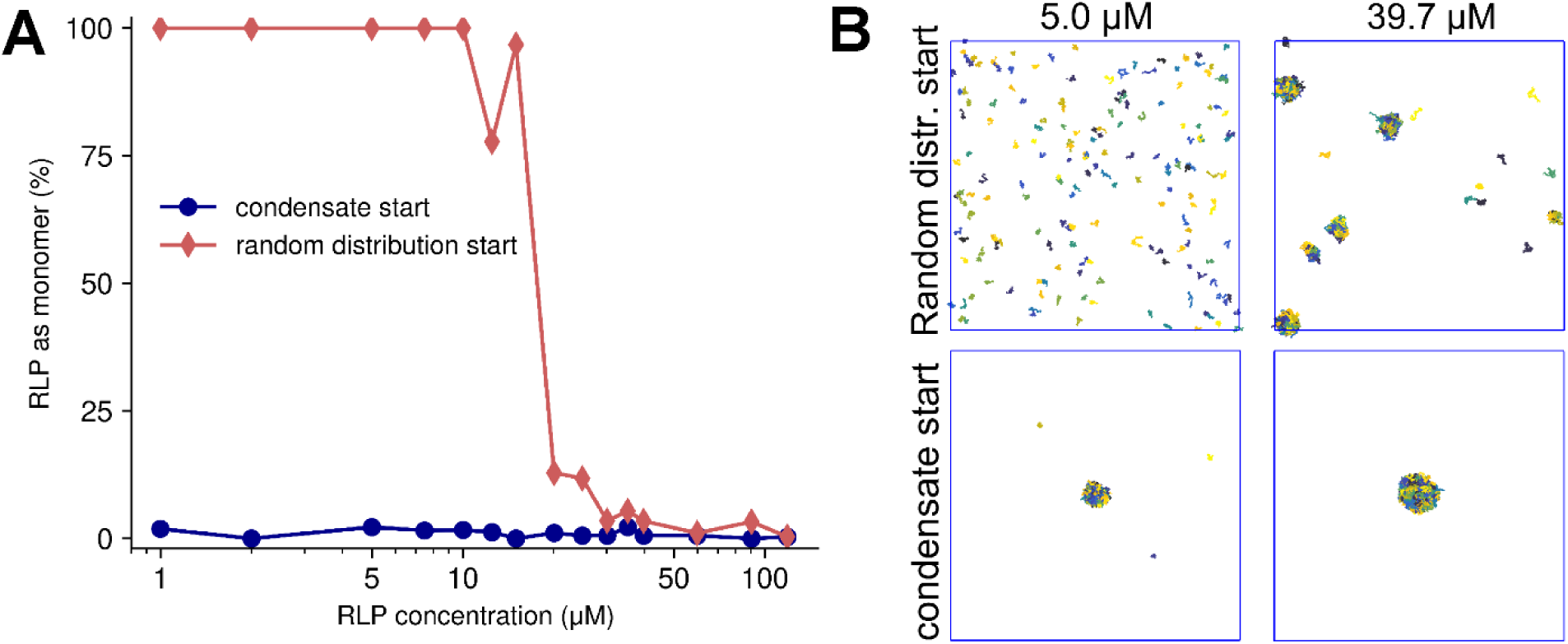
Impact of starting conditions on RLP phase separation. (A) Monomer fraction after 5 µs (averaged over the last micro-second) as a function of the initial RLP concentration initiated from 180 copies of RLP in a random distribution (red diamonds) vs. a pre-formed condensate (blue circles). (B) Terminal snapshots of simulations at 5.0 µM and 39.7 µM RLP concentrations for random distribution start (top) and condensate start (bottom).

The hysteresis effect with finite-length simulations is illustrated in **Fig. 1** for simulations of 180 molecules of Resilin-Like Polypeptide (RLP)^47,48^ at concentrations ranging from 1 to 120 µM, starting either from a random starting distribution or a pre-formed condensate. When starting with randomly distributed monomers, no stable clusters or condensates were observed below 10 µM during a 5 µs simulation (**Fig. 1A**). In contrast, simulations started from a fully condensed phase showed minimal monomer concentrations, even at 1 µM after 5 µs, with average monomer concentrations ranging from <0.01 to 0.3 µM depending on the initial concentration (**Fig. S1**).

Additionally, convergence time toward an equilibrium monomer concentration from a random distribution is strongly dependent on the total RLP concentration (**Fig. S1**), with higher concentrations leading to a faster decline in free monomers. To address these challenges, we adopted a mixed-phase starting condition, combining both condensed and solute phases at concentrations near the expected critical threshold, typically the experimental value. This approach allows the system to equilibrate more naturally, either towards lower or higher density in the dilute phase. Overall, the mixed-phase approach provides a more reliable and unbiased strategy for estimating csat, reducing the impact of kinetic challenges introduced by extreme starting conditions.

### Parameter optimization process

Optimization of COCOMO, like other similar coarse-grained force fields^19,20,36–38^, relied heavily on single-chain properties such as the radius of gyration (*R_g_*) to parameterize non-bonded interactions. Phase separation data were primarily used for validation or further refinement, since estimating *c_sat_* via simulations is significantly more challenging than the determination of single-chain properties. In contrast, the re-parameterization of COCOMO2 was based directly on experimental phase separation and solubility data across a range of protein systems, including both intrinsically disordered proteins (IDPs) and multi-domain proteins (MDPs). The selected IDPs included four systems from the original COCOMO study^26^ (FUS LCD^49,50^, LAF-1^51^, A1 LCD^52^, and hTau40-k18^53^) along with RLP ^47,48^ and alpha-synuclein (aSyn)^54^. For multi-domain proteins, we used two systems with phase-separation data (hnRNPA1 and hSUMO_hnRNPA1)^27,36^ and three single-domain proteins (MAGOH^45,55^, RefNM^45,56^, and Y14 ^45,57^) with solubility data under comparable experimental conditions.

The parameterization approach leverages the exponential relationship between the interaction energy U and *c_sat_* known from theory^33–35:^

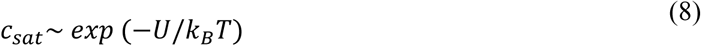

where *k_B_* is the Boltzmann constant, and T the temperature. For each of the training systems, simulations were performed with different force field parameters, keeping the number of molecules and box sizes consistent (**Table S5**). We selected simulations that achieved equilibrium between monomers in solution and those in the condensate, allowing us to estimate *C_sat_*. From a parameter set where no monomer remained in solution, we extracted ten representative conformations and used them to calculate interaction energies across different parameters. This enabled us to fit a linear relationship between *U* and the logarithm of *C_sat_*:

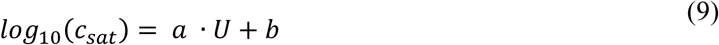

The fitting parameters a and b are reported in **Table S8**. To account for the presence of outliers, we applied the random sample consensus (RANSAC) algorithm^58^ implemented in sklearn^59^, which identifies the most reliable data points when generating linear fits.

Our parameter optimization process for COCOMO2 began with an initial scan of the parameter space using only the selected IDP systems, excluding the surface scaling factor λ as it has no impact on the IDPs (**Fig. S2**). This scan generated a preliminary set of parameters which were used as a starting point for further refinement. Next, the initial parameters were refined using the L-BFGS-B minimization algorithm^60^. This step aimed to minimize the difference between experimental *c_sat_* values and those estimated from interaction energies, improving alignment between the model and experimental data. Finally, the parameters were re-optimized using the complete set of IDPs and multi-domain proteins, this time including λ, to ensure that the force field could capture the behavior of both disordered and folded protein systems. We limited the maximum value of λ to 0.7, as that value worked well with the original COCOMO model (see below) and because there were significant changes in the morphology of the condensates with higher λ values (**Fig. S3**).

After optimizing the parameters, we ran additional simulations to evaluate the differences between experimental and estimated via *U* and simulated *c_sat_*. We used three more systems - TAP ^45,61^, GFP FUS^36,62^, WW34^45,63^ - for additional testing based on solubility and phase separation data. Additionally, we selected 14 IDPs from the original COCOMO dataset for as well as the 23 previously mentioned MDPs to test COCOMO2’s ability to reproduce radii of gyration (*R_g_*) despite not being trained against them. We further evaluated COCOMO2 performance on heterotypic protein systems (e.g., FUS LCD with (RGRGG)₅, **Table S6**) and protein-RNA systems (**Table S7**), for both of which experimental phase separation data are available.

## RESULTS

### Improvements with surface scaling for folded domains

The applicability of the COCOMO force field to proteins with folded domains was tested with a set of 23 MDPs, which were introduced previously when reparametrizing CALVADOS^64^. The primary goal was to assess how well the model could reproduce experimental *R_g_* values (**Fig. 2A**). As COCOMO is not designed to fold proteins or preserve secondary structures, an elastic network was used to preserve secondary and higher-order structural elements in folded domains, which is a common approach for coarse-grained simulations^36,65^. From 1 µs MD simulations, the original COCOMO model could not reproduce the experimental *R_g_* values, as the model consistently predicted too compact conformations, resulting in significantly underestimated *R_g_* values (purple points in **Fig. 2A**). The relative root mean-square deviation (RMSD) between experimental and calculated *R_g_* values is 36.2% (**Fig. 2B**).

**Figure 2.**
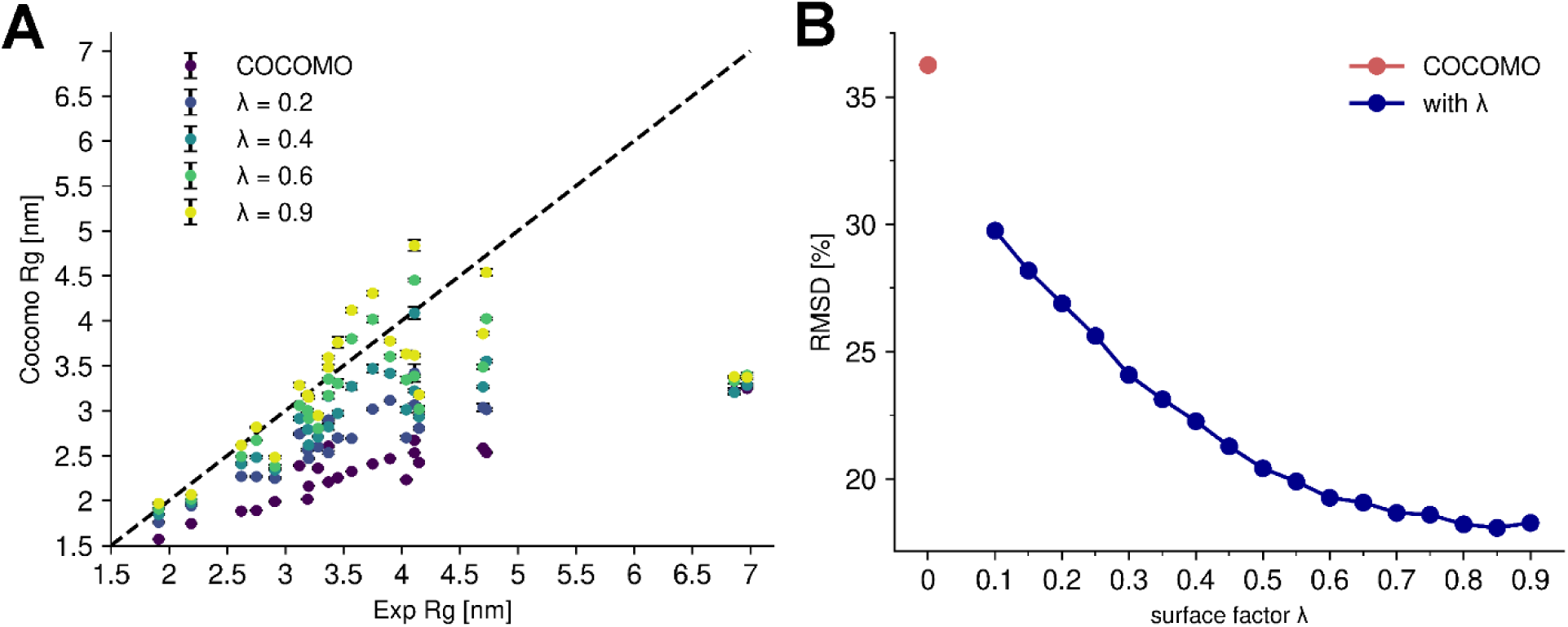
Effect of surface scaling factor (λ) on R_g_ of multi-domain proteins. (A) Comparison between the experimental radius of gyration (R_g_) of 23 multi-domain proteins and the predicted R_g_ using the original COCOMO model and COCOMO2 with varying values of λ. Without λ, COCOMO underestimates the R_g_, as indicated by the deviation from the diagonal (dashed line). (B) Relative root mean-square deviation (RMSD) between predicted and experimental R_g_ as a function of λ. The RMSD decreases as λ increases.

To correct the overestimated interactions between folded domains, we introduced a surface exposure scaling factor λ, which adjusts the interaction strength of individual residues in folded domains based on their degree of solvent accessibility. Folded proteins feature both solvent-exposed residues, which can actively participate in intermolecular interactions, and buried residues, which primarily stabilize the internal structure. Moreover, buried residues contribute less to the solvation free energy^66^. In contrast, IDPs are generally more flexible, and their residues are similarly solvent-exposed and contribute similarly to intramolecular interactions. Because COCOMO was originally parametrized for IDPs, an adjustment for buried residues in folded domains is needed to effectively account for solvation effects.

More specifically, the scaling factor λ is defined as the threshold for the ratio between a residue’s surface area in the initial structure and an amino acid-specific reference value (*S_ratio_*). If *S_ratio_* is larger than λ, the full interaction strength is applied. Otherwise, the interaction strength is scaled linearly as a function of surface exposure, reaching zero for fully buried residues (see method section, **Eq. 6**). This approach effectively reduces the inter- and intramolecular interactions of folded domains while leaving the interactions of flexible domains and IDPs unchanged. It is important to note that the folded domains are kept intact via elastic network restraints so that the degree of solvent exposure remains the same throughout a simulation of a given system with folded domains. This allowed us to determine which residues have reduced interactions once at the beginning of the simulations and then simply apply different parameters to those residues without having to introduce a surface-exposure dependent term that is costly to evaluate continuously during simulations^30,31^.

The introduction of λ significantly improved the agreement between the experimental and predicted *R_g_* values for MDPs (**Fig. 2A, B**), reducing the RMSD from 36.2% up to 18.1% for a λ of 0.85. Beyond single-chain properties, the scaling factor also affects phase-separation behavior, as illustrated with the folded protein MAGOH^45,55^ (**Fig. 3**). At low λ values, where reduced interactions mainly affect buried residues, few free monomers remain, leading to a low *c_sat_*. At λ ≈ 0.45, the predicted critical concentration is similar to the experimental solubility value of 0.11 mM. W e note that the logarithm of *c_sat_* depends linearly on λ, as expected from **Eq. 8**^33–35^ since interaction strength scales with λ and critical concentrations for phase transitions depend on interaction strength. We will take advantage of this relationship in the next section. λ-scaled interactions significantly improve *R_g_* and solubility estimates. There is also improvement in the agreement between experimental and simulated *c_sat_* for IDP systems including folded domains (**Fig. 4**), but it was difficult to identify an optimal value of λ with the original COCOMO model without further optimization.

**Figure 3.**
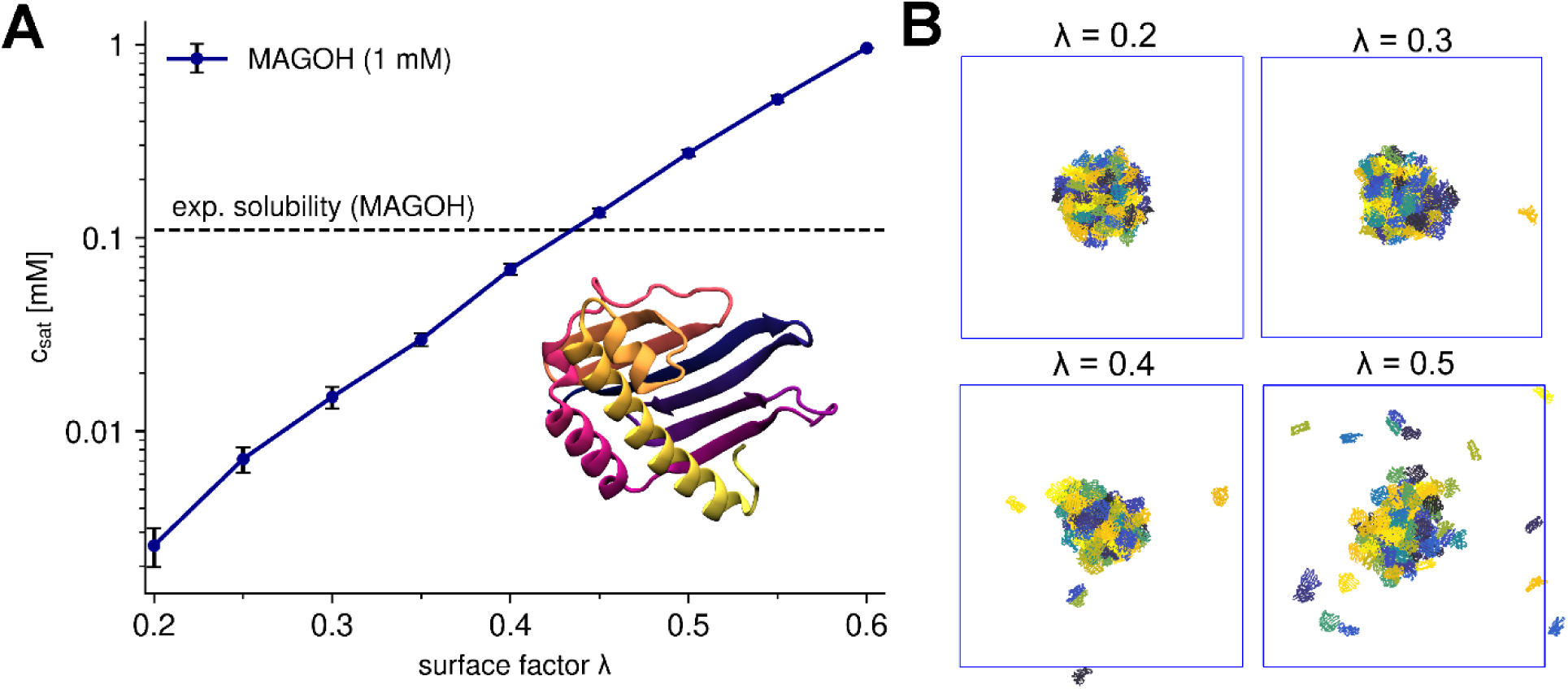
Effect of surface scaling factor (λ) on saturation concentration (c_sat_) on the example of MAGOH. (A) Effect of λ on the c_sat_ of MAGOH. The critical concentration is calculated as the average free monomer concentration over the last 5 µs in a 10 µs long simulation. As λ increases, the predicted critical concentration of MAGOH approaches the experimental solubility threshold (dashed line). (D) Representative snapshots of MAGOH simulations with different λ.

**Figure 4.**
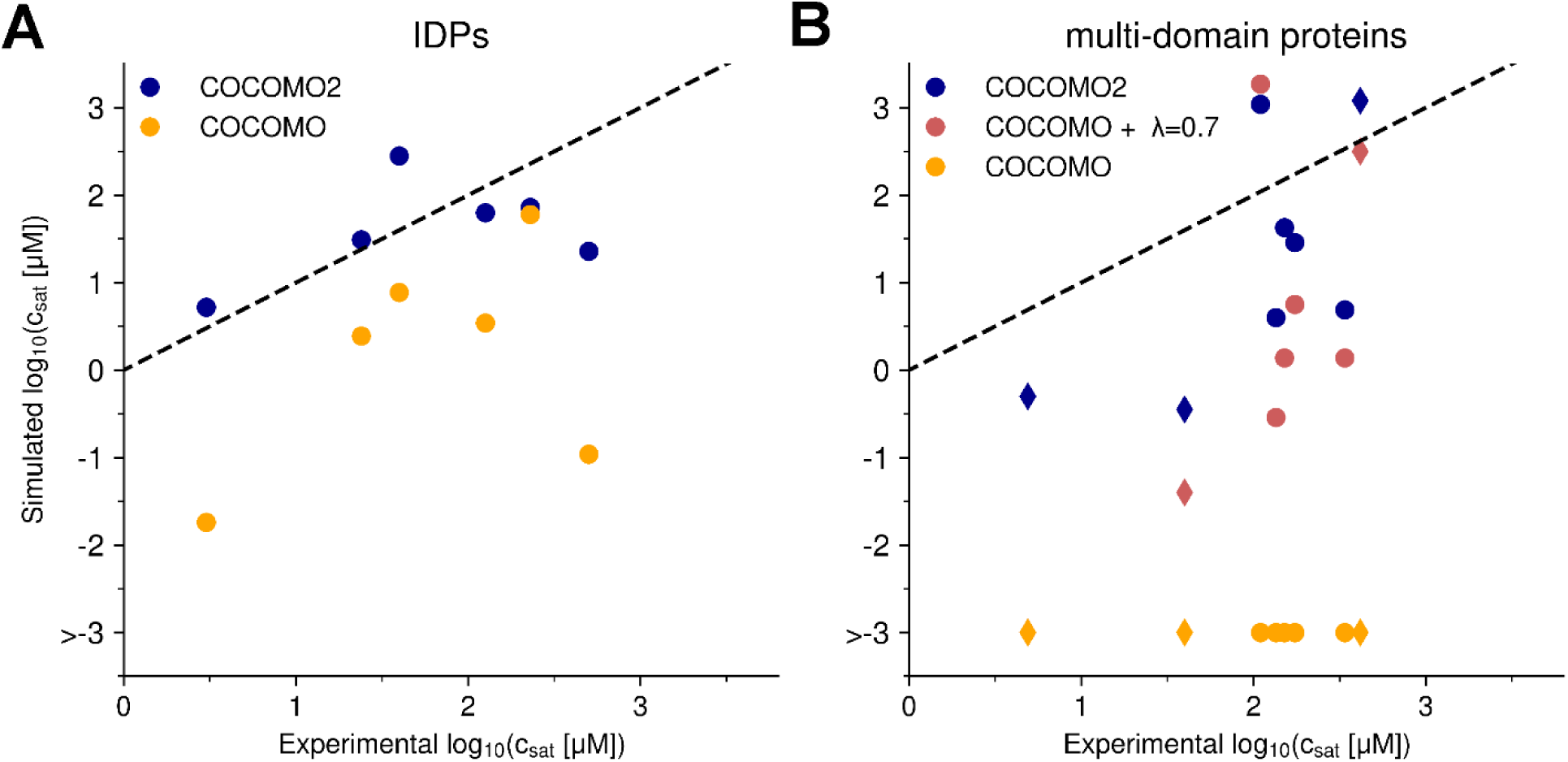
Dependency between potential energy and critical concentration for IDPs and folded Proteins. Comparison between the experimental and simulated c_sat_ values for IDPs (A) and multi-domain proteins (B) for different versions of the model: COCOMO2, COCOMO with λ=0.7, and COCOMO without surface scaling. The left panel shows results for the six IDPs, while the right panel presents data for proteins with folded domains. Circles represent the five proteins with folded domains used for parameterization, while diamonds indicate three additional systems (TAP, GFP FUS, WW34) used for validation. With the original COCOMO, we observed cases where no chains left the condensate during the simulation time and therefore c_sat_ could not be evaluated.

### Optimizing force field parameters to reproduce critical concentrations

To further optimize COCOMO, we incorporated solubility and phase separation data directly into the parameterization process. While single-chain properties, such as *R_g_*, tend to converge fast and require just one molecule, optimizing parameters based on *c_sat_* presents a greater challenge. Optimization becomes more feasible when direct correlations exist between *c_sat_* and interaction parameters or with parameter-dependent quantities that can be calculated from pre-existing structural ensembles. This avoids the need to run extensive simulations repeatedly to determine *c_sat_* empirically for each parameter set. One such useful correlation we identified is the relationship between *c_sat_* and λ.

Based on **Eq. 8**, we hypothesized that the potential energy averaged over a limited number of representative condensate structures could serve as a reliable proxy for estimating *c_sat_* (**Fig. 5**). To explore these relationships, we ran simulations for six IDPs and five folded proteins, systematically varying key parameters: the surface scaling factor λ, the Lennard-Jones potential depth (ε), and the solvation parameter (A_0_) for polar and hydrophobic residues (*cf.* Methods section). The starting configurations for these simulations were a combination of pre-formed condensates and randomly distributed monomers at concentrations close to the experimental *c_sat_*. As long as condensates did not fully dissolve or monomers did not entirely condense, we could identify meaningful correlations between force field parameters and *c_sat_*.

**Figure 5.**
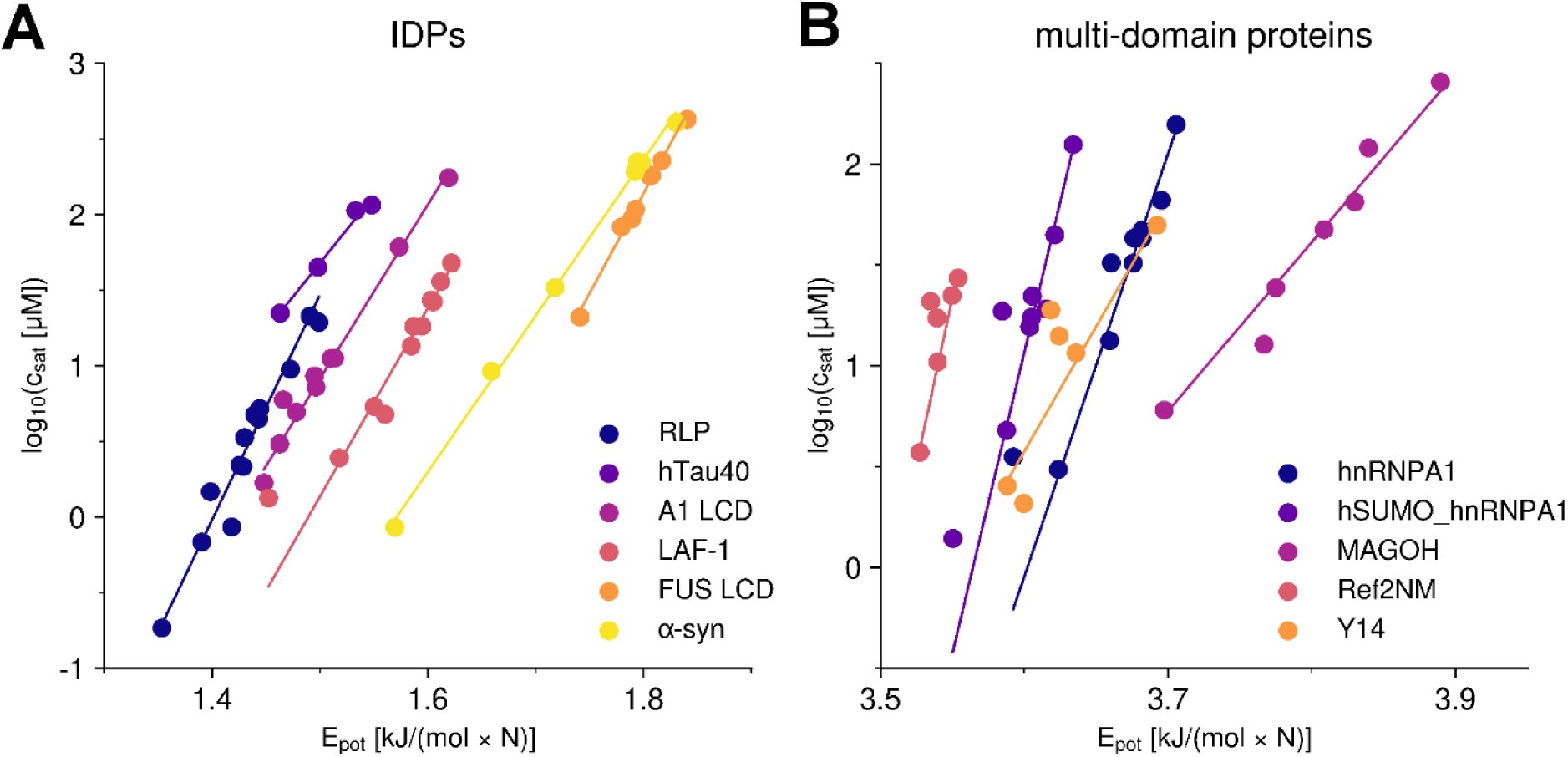
Dependency between potential energy and critical concentration for IDPs and multi-domain proteins. (A, B) Comparison between the average per-residue potential energy calculated from condensate structures and the logarithm of the critical concentration (in mM) with varying parameters in the COCOMO force field. The critical concentration was determined by averaging the monomer concentration over the final microsecond of a 5 µs simulation. (A) shows results for six IDPs, while (B) presents data for five folded proteins. For visualization purposes, the x-axis for MAGOH in (B) has been shifted by -1.9 kJ/(mol · N) . A random sample consensus (RANSAC) model was applied to determine the linear fits shown in both panels, ensuring that outliers did not skew the trendlines.

Across all tested systems, we observed linear correlations between the potential energy and the logarithm of *c_sat_*, although more outliers appeared in simulations involving multi-domain proteins (**Fig. 5B**), likely because of the structural complexity and heterogeneity of multi-domain proteins and/or incomplete sampling. To mitigate the effect of outliers, we applied the random sample consensus (RANSAC) algorithm^58^, which identifies the most reliable data points when generating linear fits.

With these linear relationships established, we optimized the force field parameters toward experimental values of *c_sat_* without requiring time-consuming, fully converged simulations. The optimization process involved three key steps: First, we performed a scan of the parameter space using only IDPs systems to generate a preliminary set of parameters expected to result in relatively low deviations from the experimental *c_sat_* values (**Fig. S2**). These initial parameter guesses were then refined through iterative minimization in the second step. Here, the goal was to minimize the difference between the experimental values and those estimated from interaction energies. Finally, we minimized the model against the experimental values by including the full set of IDPs and multi-domain proteins, incorporating λ into the optimization to ensure that the model accurately captures the behavior of both disordered and folded proteins.

To validate the optimized parameters, we ran additional simulations and compared critical concentrations estimated from potential energy with those calculated directly from the simulations using the optimized parameters (**Fig. S4, S5**). The RMSD for IDPs was low at 0.21, indicating strong predictive accuracy, with somewhat greater variability for folded proteins, given an RMSD of 0.97. The larger discrepancy between predicted and actual *c_sat_* values from simulations was most pronounced for hSUMO_hnRNPA1, likely due to a combination of multiple folded domains and disordered regions in the same system. Moreover, our approach implicitly assumes that changes in the force field do not significantly alter the morphology of the condensate, for which we have no information from the experiment. In the case of hSUMO_hnRNPA1, higher λ values led to more surface-exposed folded domains (**Fig. S3**). Because of that, we limited the maximum value of λ to 0.7 during our optimization.

COCOMO2 demonstrated substantially improved predictive performance compared to the original COCOMO model with λ (**Fig. 4**), reducing the RMSD between simulated and experimental critical concentrations from 1.94 to 0.68 for IDPs and from 2.34 to 1.28 for partially folded proteins. Without surface scaling, the original COCOMO model clearly produced overly stable condensates across all multi-domain systems tested, as there was no coexistence with monomers in the dilute phase (**Fig. 4**).

Overall, our optimization strategy provided a computationally efficient approach to incorporating phase separation and solubility data, resulting in improved predictive accuracy As a result, we expect COCOMO2 to be more versatile in modeling both disordered and folded proteins.

### Further evaluation of COCOMO2

We further evaluated COCOMO2 on systems not used for optimization. We tested COCOMO2 performance for predicting *R_g_* values across both IDPs and proteins with folded domains (**Fig. 6**). As mentioned before, experimental *R_g_* values were not used to optimize COCOMO2. We found notable improvement with COCOMO2. For the IDPs, we selected a representative sub-section of systems tested in the original COCOMO paper^26^ to cover a wide range of experimental *R_g_* values. The original COCOMO model performed well for IDPs with up to ∼150 amino acid residues but underestimated the *R_g_* values of longer chains. With the COCOMO2 parameters the interaction energy is reduced, allowing for a more extended conformational ensemble, especially for longer chains. As a result, there is marked improvement when comparing predicted *R_g_* values with experimental data, reducing χ^2^ from 3.65 to 2.11 across the IDP set (**Fig. 6B**). While improvements were observed across all tested systems, the effects were especially notable for MDPs. The original COCOMO parameters consistently underestimated the *R_g_* values. The introduction of the surface scaling (λ = 0.7) already provided a substantial improvement by decreasing the interactions of buried residues, reducing the χ^2^ from 0.75 to 0.55. With the reparametrized COCOMO2 model, there was further reduction of χ^2^ to 0.24, with a marked impact on the system with extended disordered regions and small folded domains like HeV_V and NiV_V^67^, whose *R_g_* values are only weakly affected by varying λ values. As such, despite not being explicitly parameterized against *R_g_* values, COCOMO2 provides a substantial improvement over the original COCOMO in accurately predicting *R_g_* across a wide range of systems.

**Figure 6.**
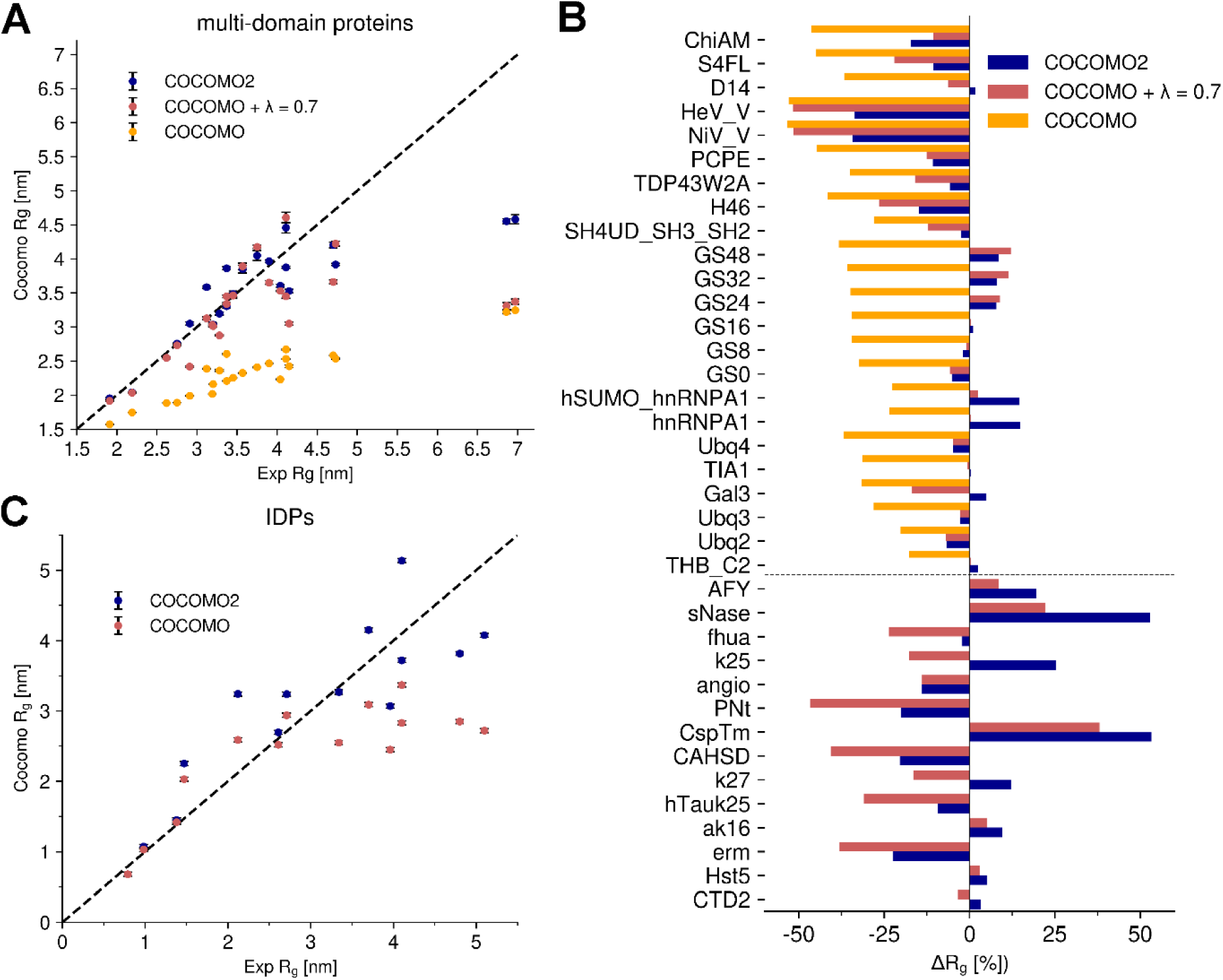
Comparison of experimental vs simulated Radius of Gyration (R_g_) for IDPs and multi-domain proteins. (A) Experimental vs simulated R_g_ for folded proteins using COCOMO2, COCOMO with λ=0.7, and the original COCOMO parameters. (B) Comparison between experimental and simulated Rg for IDPs using COCOMO2 and the original COCOMO parameters (λ does not affect IDPs and disordered regions). COCOMO2 provides better agreement with experimental R_g_ values than the COCOMO model, especially for systems with more residues. (C) Relative differences in R_g_ (ΔR_g_ [%]) between simulation and experimental data for all tested systems, grouped by protein type (top folded proteins, bottom IDPs). Negative values indicate underestimation of R_g_ by the model, while positive values indicate overestimation. COCOMO2 exhibits smaller deviations compared to the other models, with the largest improvements observed for folded proteins.

We also observed improved estimates of *c_sat_* for three additional folded systems that were not included in the parameterization (decrease in RMSD from 2.75 to 1.34, **Fig. 4**), demonstrating that the improved performance of COCOMO2 is not limited to the training set. As an additional test, we evaluated whether COCOMO2 could still accurately predict heterotypic phase separation^50^ and protein-RNA phase separation as with the original COCOMO model. We set up simulations with 200 µM FUS LCD, partially as monomers, partially in a pre-formed condensate, together with different ratios of (RGRGG)_5_ peptides (**Fig. 7A**). This system was used as a test set in the original COCOMO paper^26^ due to the availability of heterotypic phase separation data. We found that at 1:1 and 1:2 FUS LCD/(RGRGG)_5_ ratios, the condensate dissociates, while it remained stable at ratios of 1:5 and 1:10 after ten µs simulation. This result suggests that higher concentrations of (RGRGG)_5_ are required to stabilize the heterotypic condensate, consistent with the experimental results^50^, confirming that COCOMO2 retains the ability to capture the system-dependence determinants of heterotypic phase separation.

**Figure 7.**
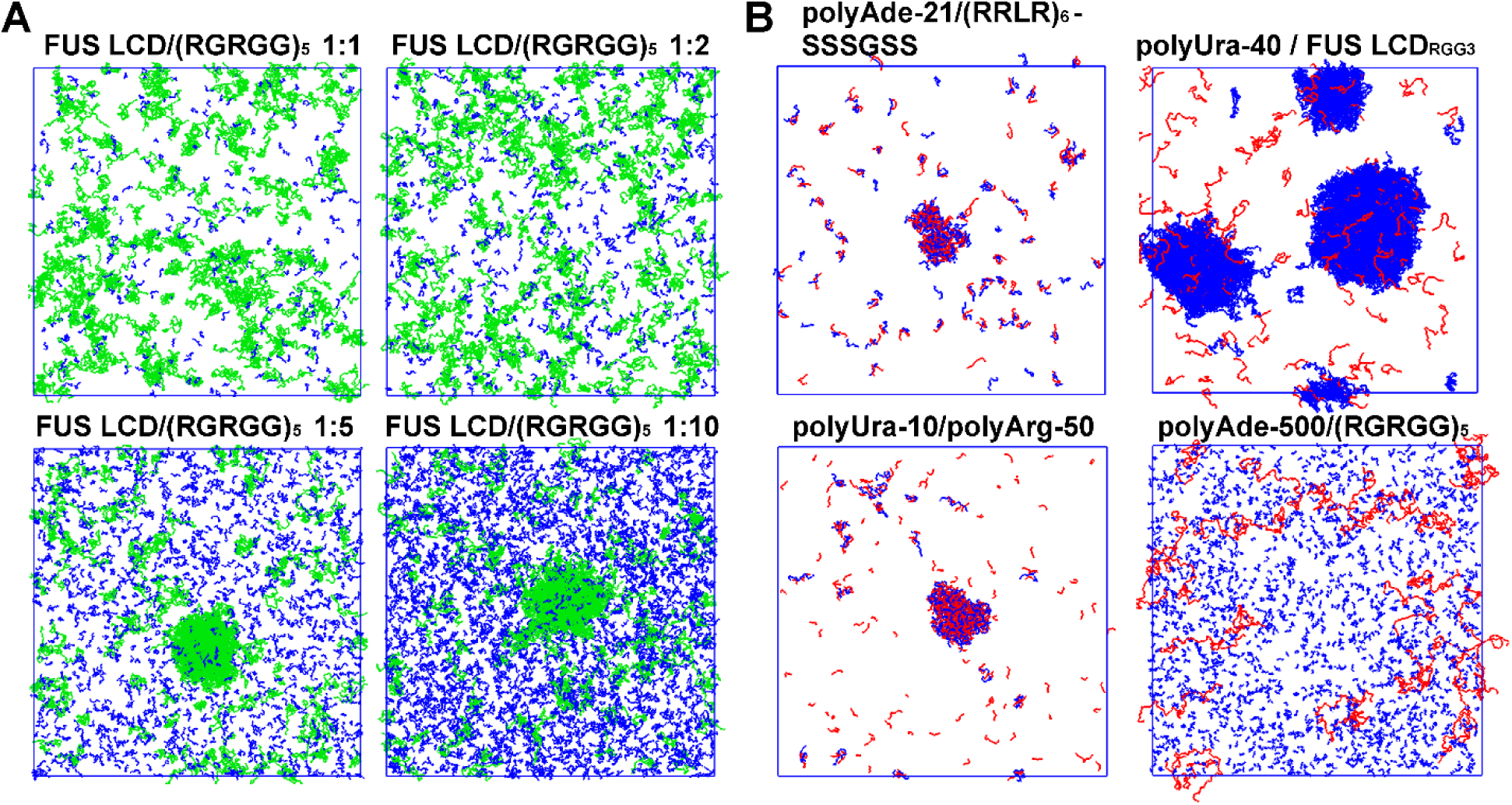
Protein heterotypic and protein-RNA phase separation. (A) Simulation snapshots after 10 µs simulations of mixes between FUS LCD (green) and (RGRGG)₅ (blue) at varying concentration ratios. All simulations were set up with 200 µM FUS LCD concentration with a pre-formed condensate already present. For low (RGRGG)₅ concentrations, the condensate has dissolved. (B) Simulation snapshots for various RNA-protein systems at concentration with experimentally observed phase separation. Proteins are colored in blue, and RNA molecules in red. Systems include polyAde-21/(RRLR)_6_, polyUra-40/FUS LCD_RGG3_, polyUra-10/polyArg-50, and polyAde-500/(RGRGG)₅ with a randomly dispersed initial state. The final states show varying degrees of phase separation, with polyAde-500/(RGRGG)₅ remaining largely dispersed.

Additionally, we tested four RNA-IDP systems (polyAde-21/(RRLR)_6_^68^, polyUra-40/FUS LCD ^50^, polyUra-10/polyArg-50^69^, and polyAde-500/(RGRGG)₅^70^) at concentrations where phase separation was observed experimentally. These simulations started from a randomly distributed state. We observed phase separation in three of the four systems, with polyAde- 500/(RGRGG)₅ remaining dispersed. Compared to the original COCOMO simulations, a higher number of IDP molecules remained in the solution, consistent with weaker protein interactions in the revised force field. Therefore, COCOMO2 agrees qualitatively with the experiment for three out of the four systems without further optimization of the protein-RNA interactions. Further optimization of the protein-RNA interactions may be a subject of future work but may require more extensive experimental data, in particular, critical concentrations of peptides and nucleic acids, in addition to qualitative observations of phase separation at certain concentrations.

## DISCUSSION and CONCLUSION

We present an updated version of the COCOMO coarse-grained model for peptides and proteins that extends its application to folded proteins and IDPs with folded domains. Simulations of folded proteins with COCOMO requires elastic network model restraints to keep secondary and tertiary structures intact but opens up new applications, including the simulation of complex assembly processes.

One of the core enhancements in COCOMO2 is the introduction of surface scaling factor λ, which adjusts the interaction strength of residues based on solvent accessibility. This modification addresses a key limitation of the original model, which overestimated the interaction strength of folded proteins. In COCOMO2, buried residues in folded domains now contribute less to intermolecular interactions, ensuring a more accurate representation of folded domains. However, this change alone was insufficient to fully resolve the underestimation of critical concentrations observed for both IDPs and multi-domain proteins.

We further improved the model by incorporating phase-separation data directly into parameterization, leveraging a linear relationship between potential energy in condensates and the logarithm of critical concentration. Until now, most similar, single-bead-per-residue coarse-grained models for IDPs relied primarily on single-chain properties like the radius of gyration to derive the force field parameters for parameterization and used phase-separation data as control or validation. Our approach could potentially also benefit other force fields. We observed that this method performs better for IDPs than for multi-domain proteins, where deviations between simulated and experimental critical concentrations were more pronounced. This suggests that further optimization via conventional cycles of parameter variations and trial simulations could be possible, but the greatly improved accuracy of COCOMO2 may already be sufficient for typical applications that are focused on qualitative or semi-quantitative predictions of interaction preferences and the resulting condensation and/or clustering of proteins including folded domains.

Among other available methods, the recently updated CALVADOS3^36^ model predicts *c_sat_* with similar deviation for multi-domain proteins as COCOMO2, typically about one order of magnitude or more different from experimental values. However, COCOMO2 and CALVADOS3 adopt different design philosophies: CALVADOS3 extends its applicability from IDPs to folded proteins by shifting the bead location in folded domains from the Cα position to the center of mass which may implicitly capture increased interactions of surface-exposed residues. Instead, COCOMO2 uses a surface scaling approach that explicitly considers residue burial. Moreover, COCOMO2 emphasizes a minimal number of parameters by grouping polar and hydrophobic residues, while CALVADOS3 assigns individual parameters to each amino acid. As a result, CALVADOS3 is more sensitive to the exact amino acid sequence of a given protein and can capture the effects of mutations better than COCOMO2, but the higher level of detail may limit generalizability. Whether this is, in fact, a concern remains to be seen, given the relatively small amount of experimental data on critical concentrations for phase separation available for comparison.

The optimization of COCOMO2 using phase-separation also improved the agreement with experimental *R_g_* values for both IDPs and multidomain properties. The original COCOMO worked well for smaller IDPs but underestimated the *R_g_* of longer chains. In COCOMO2, weaker interaction parameters for polar and hydrophobic residues allow for more expanded conformations, improving accuracy. However, comparing again with other methods, CALVADOS3 provides significantly more accurate *R_g_* predictions since it was explicitly optimized to match experimental *R_g_* values.

COCOMO2 parameterization and validation focused on critical concentrations for phase separation and single-chain *R_g_* values, as such data is readily available from experiments. However, we expect that COCOMO2 will also be useful for the study of specific assembly processes. In assembly processes, it is important to avoid non-specific aggregation but favor interactions at specific sites that lead to organized higher-order structures. We expect that the generic nature of COCOMO2 is well-suited to address the avoidance of non-specific aggregation and that preference for interactions at specific sites can be added via knowledge-based potentials^71,72^. The advantage of such an approach compared to more coarse-grained models used previously for studying assembly processes^73–75^ is that the residue-level model of COCOMO provides a higher level of detail with a path to connect with atomistic models^76^ in multi-scale applications. These strategies will be explored in future studies.

In conclusion, COCOMO2 offers a comprehensive framework for modeling interactions between peptides and proteins and nucleic acids that is extended to also include folded proteins. Its balance between simplicity and physical accuracy positions COCOMO2 as a valuable resource for understanding biomolecular condensates and complex molecular environments. The highly efficient model makes it possible to study phase separation and assembly processes on sub-µm scales and millisecond time scales.

## ACKNOWLEDGEMENTS

Research was primarily supported as part of the Center for Catalysis in Biomimetic Confinement, an Energy Frontier Research Center funded by the U.S. Department of Energy, Office of Science, Basic Energy Sciences under Award #DE-SC0023395. In addition, Lisa Lapidus and Michael Feig acknowledge support by the National Science Foundation under Award #MCB1817307, (prediction of concentration-dependent liquid-liquid phase separation) and Michael Feig acknowledges support by the National Institutes of Health (NIGMS) under Award #R35GM126948, (development of a coarse-grained model for use in multi-scale applications of cellular environments).

