## Supplementary Material for "COCOMO2: A coarse-grained model for interacting folded and disordered proteins"

**Tables S1-S9**

**Figures S1-S5**

**Supplementary References**

**Table S1. Amino acid-specific reference surface areas ($S_{ref}$) used in COCOMO2.**

| Residue | $S_{ref}$ [nm$^2$] |
|---|---|
| ALA | 0.796 |
| ARG | 1.921 |
| ASN | 1.281 |
| ASP | 1.162 |
| CYS | 1.074 |
| GLN | 1.575 |
| GLU | 1.462 |
| GLY | 0.544 |
| HIS | 1.634 |
| ILE | 1.410 |
| LEU | 1.519 |
| LYS | 1.923 |
| MET | 1.620 |
| PHE | 1.869 |
| PRO | 0.974 |
| SER | 0.933 |
| THR | 1.128 |
| TRP | 2.227 |
| TYR | 2.018 |
| VAL | 1.232 |

**Table S2. Multi-domain proteins used for testing of COCOMO2 and the effect of λ.**

| Protein | Rg [nm] | N | Folded domains | Ions [M] | pH | Reference |
|---|---|---|---|---|---|---|
| THB_C2 | 1.91 | 137 | [6, 42] [50, 137] | 0.15 | 6.5 | Michie *et al.* [1] |
| Ubq2 | 2.19 | 162 | [11, 82] [87, 158] | 0.33 | 8.0 | Jussupow *et al.*[2] |
| Ubq3 | 2.62 | 228 | [1, 72] [77, 148] [153, 224] | 0.33 | 8.0 | Jussupow *et al.*[2] |
| Gal3 | 2.91 | 250 | [117, 250] | 0.04 | 7.0 | Lin *et al.*[3] |
| TIA1 | 2.75 | 275 | [6, 82] [95, 172] [190, 275] | 0.10 | 6.0 | Sonntag *et al.*[4] |
| Ubq4 | 3.19 | 304 | [1, 72] [77, 148] [153, 224] [229, 300] | 0.33 | 8.0 | Jussupow *et al.*[2] |
| hnRNPA1 | 3.12 | 314 | [11, 89] [105, 179] | 0.15 | 7.5 | Martin *et al.*[5] |
| hSUMO_hnRNPA1 | 3.37 | 433 | [44, 114] [132, 209] [224, 298] | 0.10 | 7.5 | Martin *et al.*[5] |
| GS0 | 3.2 | 470 | [1, 226] [256, 470] | 0.15 | 7.4 | Moses *et al.*[6] |
| GS8 | 3.37 | 486 | [1, 226] [272, 486] | 0.15 | 7.4 | Moses *et al.*[6] |
| GS16 | 3.45 | 502 | [1, 226] [288, 502] | 0.15 | 7.4 | Moses *et al.*[6] |
| GS24 | 3.57 | 518 | [1, 226] [304, 518] | 0.15 | 7.4 | Moses *et al.*[6] |
| GS32 | 3.75 | 534 | [1, 226] [320, 534] | 0.15 | 7.4 | Moses *et al.*[6] |
| GS48 | 4.11 | 566 | [1, 226] [352, 566] | 0.15 | 7.4 | Moses *et al.*[6] |
| SH4UD_SH3_SH2 | 3.28 | 264 | [94, 150] [166, 258] | 0.216 | 8.0 | Gurumoorty *et al.*[7] |
| H46 | 4.15 | 381 | [140, 355] | 0.163 | 6.5 | Elena-Real *et al.*[8] |
| TDP43W2A | 4.11 | 415 | [5, 77] [107, 177] [193, 260] [321, 329] | 0.312 | 8.0 | Wright *et al.*[9] |
| PCPE | 4.04 | 424 | [12, 125] [134, 249] [293, 412] | 0.506 | 7.4 | Bernocco *et al.*[10] |
| NiV_V | 6.97 | 457 | [406, 457] | 0.232 | 8.0 | Salladini *et al.*[11] |
| HeV_V | 6.86 | 458 | [404, 456] | 0.232 | 8.0 | Salladini *et al.*[11] |
| D14 | 3.9 | 483 | [31, 121] [157, 246] [265, 354] [400, 479] | 0.156 | 7.5 | Hajizadeh *et al.*[12] |
| S4FL | 4.7 | 552 | [15, 138] [287 542] | 0.169 | 7.2 | Gomes *et al.*[13] |
| ChiAM | 4.73 | 682 | [8, 89] [92, 172] [178, 257] [266, 356] [359, 462] [471, 567] [578, 668] | 0.282 | 8.0 | Mazurkewich *et al.*[14] |

**Table S3. RLP phase separation hysteresis simulation set.**

| Protein | $N_{chain}$ (cond) | $N_{chain}$ (free) | c [µM] | Box [nm] |
|---|---|---|---|---|
| RLP | 0 | 180 | 1.0 | 669 |
| RLP | 0 | 180 | 2.0 | 531 |
| RLP | 0 | 180 | 5.0 | 391 |
| RLP | 0 | 180 | 7.5 | 342 |
| RLP | 0 | 180 | 10.0 | 310 |
| RLP | 0 | 180 | 12.5 | 288 |
| RLP | 0 | 180 | 15.0 | 271 |
| RLP | 0 | 180 | 20.1 | 246 |
| RLP | 0 | 180 | 24.9 | 229 |
| RLP | 0 | 180 | 30.1 | 215 |
| RLP | 0 | 180 | 35.2 | 204 |
| RLP | 0 | 180 | 39.7 | 196 |
| RLP | 0 | 180 | 59.8 | 171 |
| RLP | 0 | 180 | 90.4 | 149 |
| RLP | 0 | 180 | 118.8 | 136 |
| RLP | 180 | 0 | 1.0 | 669 |
| RLP | 180 | 0 | 2.0 | 531 |
| RLP | 180 | 0 | 5.0 | 391 |
| RLP | 180 | 0 | 7.5 | 342 |
| RLP | 180 | 0 | 10.0 | 310 |
| RLP | 180 | 0 | 12.5 | 288 |
| RLP | 180 | 0 | 15.0 | 271 |
| RLP | 180 | 0 | 20.1 | 246 |
| RLP | 180 | 0 | 24.9 | 229 |
| RLP | 180 | 0 | 30.1 | 215 |
| RLP | 180 | 0 | 35.2 | 204 |
| RLP | 180 | 0 | 39.7 | 196 |
| RLP | 180 | 0 | 59.8 | 171 |
| RLP | 180 | 0 | 90.4 | 149 |
| RLP | 180 | 0 | 118.8 | 136 |

**Table S4. Homotypic protein phase separation systems for IDPs and multi-domain proteins.**

| Protein | $c_{sat,\,exp}$ [µM] | $N_{chain}$ (cond) | $N_{chain}$ (free) | Box [nm] | Folded domains | Reference |
|---|---|---|---|---|---|---|
| hTau40-k18 | 40 | 229 | 27 | 104 | | Ambadipudi *et al.* [15] |
| A1 LCD | 125 | 217 | 77 | 101 | | Bremer *et al.*[16] |
| LAF1 | 24 | 178 | 81 | 177 | | Elbaum-Garfinkle *et al.*[17] |
| RLP | 3 | 180 | 15 | 171 | | Dai *et al.*[18] |
| FUS LCD | 235 | 172 | 35 | 69 | | Kaur *et al.*[19] |
| αSyn | 500 | 0 | 300 | 100 | | Ray *et al.*[20] |
| hnRNPA1 | 173 | 200 | 77 | 90 | [11, 89] [105, 179] | Martin *et al.*[5] |
| hSUMO_hnRNPA1 | 136 | 200 | 60 | 90 | [44, 114] [132, 209] [224, 298] | Martin *et al.*[5] |
| MAGOH | 110 | 100 | 2 | 50 | [1 146] | Golovanov *et al.*[21] |
| Ref2NM | 150 | 170 | 30 | 80 | [87 166] | Golovanov *et al.*[21] |
| Y14 | 340 | 142 | 58 | 63 | [71 149] | Golovanov *et al.*[21] |
| TAP | 40 | 69 | 31 | 100 | [118 194] [204 355] [380 548] [564 619] | Golovanov *et al.*[21] |
| GFP FUS | 4.9 | 100 | 14 | 168 | [286 368] [423 451] [529 755] | Wang *et al.*[22] |
| WW34 | 420 | 400 | 600 | 85 | [15 40] [59 82] | Golovanov *et al.*[21] |

**Table S5. Parameter sets used to establish the link between potential energy and saturation concentration for IDPs and multi-domain proteins.**

| Protein | $\varepsilon_{polar}$ | $\varepsilon_{charged}$ | $\varepsilon_{hydrophobic}$ | $A_{0,polar}$ | $A_{0,hydrophobic}$ | λ | Epot [kL/(mol · N)] | $\mathrm{Log_{10}}(c_{sat})$ |
|---|---|---|---|---|---|---|---|---|
| hTau40-k18 | 0.40 | | 0.50 | 0.07 | 0 | | 1.548 | 2.063 |
| hTau40-k18 | 0.40 | | 0.41 | 0.052 | 0 | | 1.463 | 1.348 |
| hTau40-k18 | 0.40 | | 0.41 | 0.054 | 0 | | 1.498 | 1.652 |
| hTau40-k18 | 0.40 | | 0.41 | 0.056 | 0 | | 1.533 | 2.027 |
| A1 LCD | 0.40 | | 0.41 | 0.05 | 0 | | 1.448 | 0.225 |
| A1 LCD | 0.40 | 0.35 | 0.41 | 0.05 | 0 | | 1.462 | 0.484 |
| A1 LCD | 0.40 | 0.30 | 0.41 | 0.05 | 0 | | 1.478 | 0.694 |
| A1 LCD | 0.40 | 0.25 | 0.41 | 0.05 | 0 | | 1.495 | 0.933 |
| A1 LCD | 0.40 | 0.20 | 0.41 | 0.05 | 0 | | 1.514 | 1.051 |
| A1 LCD | 0.375 | | 0.41 | 0.05 | 0 | | 1.51 | 1.048 |
| A1 LCD | 0.35 | | 0.41 | 0.05 | 0 | | 1.573 | 1.786 |
| A1 LCD | 0.40 | | 0.50 | 0.08 | 0 | | 1.619 | 2.243 |
| A1 LCD | 0.40 | | 0.50 | 0.07 | 0 | | 1.466 | 0.776 |
| A1 LCD | 0.38 | 0.325 | 0.41 | 0.05 | 0 | | 1.496 | 0.86 |
| LAF1 | 0.40 | | 0.50 | 0.06 | 0 | | 1.452 | 0.125 |
| LAF1 | 0.40 | | 0.50 | 0.07 | 0 | | 1.612 | 1.556 |
| LAF1 | 0.385 | 0.325 | 0.41 | 0.05 | 0 | | 1.594 | 1.262 |
| LAF1 | 0.38 | 0.325 | 0.41 | 0.05 | 0 | | 1.605 | 1.423 |
| LAF1 | 0.40 | | 0.41 | 0.05 | 0 | | 1.518 | 0.392 |
| LAF1 | 0.40 | 0.35 | 0.41 | 0.05 | 0 | | 1.55 | 0.731 |
| LAF1 | 0.40 | 0.30 | 0.41 | 0.05 | 0 | | 1.585 | 1.132 |
| LAF1 | 0.40 | 0.25 | 0.41 | 0.05 | 0 | | 1.622 | 1.68 |
| LAF1 | 0.375 | | 0.41 | 0.05 | 0 | | 1.587 | 1.261 |
| LAF1 | 0.40 | 0.3875 | 0.41 | 0.05 | 0 | | 1.56 | 0.678 |
| LAF1 | 0.40 | 0.375 | 0.41 | 0.05 | 0 | | 1.603 | 1.434 |
| RLP | 0.3875 | 0.3875 | 0.41 | 0.05 | 0 | | 1.398 | 0.167 |
| RLP | 0.375 | 0.375 | 0.41 | 0.05 | 0 | | 1.444 | 0.718 |
| RLP | 0.3625 | 0.3625 | 0.41 | 0.05 | 0 | | 1.49 | 1.329 |
| RLP | 0.40 | | 0.41 | 0.05 | 0 | | 1.353 | -0.734 |
| RLP | 0.40 | 0.35 | 0.41 | 0.05 | 0 | | 1.39 | -0.165 |
| RLP | 0.40 | 0.30 | 0.41 | 0.05 | 0 | | 1.43 | 0.525 |
| RLP | 0.40 | 0.25 | 0.41 | 0.05 | 0 | | 1.473 | 0.977 |
| RLP | 0.375 | | 0.41 | 0.05 | 0 | | 1.426 | 0.343 |
| RLP | 0.35 | | 0.41 | 0.05 | 0 | | 1.499 | 1.287 |
| RLP | 0.39 | 0.325 | 0.41 | 0.05 | 0 | | 1.418 | -0.065 |
| RLP | 0.38 | 0.325 | 0.41 | 0.05 | 0 | | 1.429 | 0.334 |
| RLP | 0.37 | 0.325 | 0.41 | 0.05 | 0 | | 1.439 | 0.675 |
| RLP | 0.40 | | 0.50 | 0.05 | 0 | | 1.443 | 0.65 |
| RLP | 0.40 | | 0.50 | 0.06 | 0 | | 1.398 | 0.167 |
| RLP | 0.40 | | 0.50 | 0.07 | 0 | | 1.444 | 0.718 |
| FUS LCD | 0.40 | | 0.50 | 0.06 | 0 | | 1.741 | 1.322 |
| FUS LCD | 0.39 | 0.325 | 0.41 | 0.05 | 0 | | 1.789 | 1.971 |
| FUS LCD | 0.38 | 0.325 | 0.41 | 0.05 | 0 | | 1.817 | 2.356 |
| FUS LCD | 0.40 | 0.3875 | 0.41 | 0.05 | 0 | | 1.808 | 2.259 |
| FUS LCD | 0.40 | 0.36 | 0.41/0.47* | 0.05 | 0 | | 1.779 | 1.917 |
| FUS LCD | 0.40 | 0.32 | 0.41/0.53* | 0.05 | 0 | | 1.793 | 2.034 |
| FUS LCD | 0.40 | | 0.41 | 0.052 | 0 | | 1.806 | 2.255 |
| FUS LCD | 0.40 | | 0.41 | 0.054 | 0 | | 1.841 | 2.63 |
| αSyn | 0.20 | | 0.443 | 0.025 | 0.0098 | | 1.792 | 2.285 |
| αSyn | 0.238 | | 0.45 | 0.022 | 0.0075 | | 1.569 | -0.067 |
| αSyn | 0.175 | | 0.443 | 0.022 | 0.0098 | | 1.83 | 2.606 |
| αSyn | 0.238 | | 0.38 | 0.022 | 0.0098 | | 1.794 | 2.348 |
| αSyn | 0.238 | | 0.443 | 0.027 | 0.015 | | 1.798 | 2.346 |
| αSyn | 0.22 | | 0.43 | 0.022 | 0.0098 | | 1.718 | 1.517 |
| αSyn | 0.238 | | 0.443 | 0.024 | 0.098 | | 1.659 | 0.964 |
| hnRNPA1 | 0.35 | | 0.41 | 0.05 | 0 | 0.50 | 3.695 | 1.821 |
| hnRNPA1 | 0.35 | | 0.41 | 0.05 | 0 | 0.40 | 3.661 | 1.511 |
| hnRNPA1 | 0.39 | 0.325 | 0.41 | 0.05 | 0 | 0.40 | 3.624 | 0.486 |
| hnRNPA1 | 0.39 | 0.325 | 0.41 | 0.05 | 0 | 0.50 | 3.659 | 1.125 |
| hnRNPA1 | 0.40 | 0.30 | 0.41 | 0.053 | 0 | 0.40 | 3.676 | 1.633 |
| hnRNPA1 | 0.40 | 0.30 | 0.41 | 0.053 | 0 | 0.30 | 3.592 | 0.549 |
| hnRNPA1 | 0.40 | 0.30 | 0.41 | 0.053 | 0 | 0.20 | 3.706 | 2.196 |

| | | | | | | | | |
|---|---|---|---|---|---|---|---|---|
| hnRNPA1 | 0.238 | | 0.442 | 0.022 | 0.098 | 0.60 | 3.682 | 1.672 |
| hnRNPA1 | 0.238 | | 0.443 | 0.022 | 0.098 | 0.60 | 3.682 | 1.631 |
| hnRNPA1 | 0.216 | | 0.479 | 0.031 | 0.0014 | 0.51 | 3.676 | 1.509 |
| hSUMO_hnRNPA1 | 0.35 | | 0.41 | 0.05 | 0 | 0.50 | 3.621 | 1.649 |
| hSUMO_hnRNPA1 | 0.35 | | 0.41 | 0.05 | 0 | 0.40 | 3.585 | 1.271 |
| hSUMO_hnRNPA1 | 0.39 | 0.325 | 0.41 | 0.05 | 0 | 0.40 | 3.55 | 0.144 |
| hSUMO_hnRNPA1 | 0.39 | 0.325 | 0.41 | 0.05 | 0 | 0.50 | 3.588 | 0.68 |
| hSUMO_hnRNPA1 | 0.40 | 0.30 | 0.41 | 0.053 | 0 | 0.40 | 3.606 | 1.346 |
| hSUMO_hnRNPA1 | 0.40 | 0.30 | 0.41 | 0.053 | 0 | 0.30 | 3.634 | 2.097 |
| hSUMO_hnRNPA1 | 0.40 | 0.30 | 0.41 | 0.053 | 0 | 0.20 | 3.604 | 1.194 |
| hSUMO_hnRNPA1 | 0.238 | | 0.443 | 0.022 | 0.098 | 0.60 | 3.615 | 1.282 |
| hSUMO_hnRNPA1 | 0.216 | | 0.479 | 0.031 | 0.0014 | 0.51 | 3.605 | 1.24 |
| hSUMO_hnRNPA1 | 0.238 | | 0.442 | 0.022 | 0.098 | 0.51 | 5.74 | 1.649 |
| MAGOH | 0.39 | 0.325 | 0.41 | 0.05 | 0 | 0.30 | 5.709 | 2.08 |
| MAGOH | 0.39 | 0.325 | 0.41 | 0.05 | 0 | 0.25 | 5.675 | 1.676 |
| MAGOH | 0.39 | 0.325 | 0.41 | 0.05 | 0 | 0.20 | 5.597 | 1.388 |
| MAGOH | 0.40 | | 0.41 | 0.05 | 0 | 0.20 | 5.667 | 0.78 |
| MAGOH | 0.40 | | 0.41 | 0.05 | 0 | 0.30 | 5.73 | 1.107 |
| MAGOH | 0.40 | | 0.41 | 0.05 | 0 | 0.40 | 5.79 | 1.813 |
| MAGOH | 0.40 | | 0.41 | 0.05 | 0 | 0.50 | 5.74 | 2.407 |
| Ref2NM | 0.40 | | 0.408 | 0.041 | 0.013 | 0.70 | 3.54 | 1.019 |
| Ref2NM | 0.216 | | 0.479 | 0.031 | 0.0014 | 0.51 | 3.554 | 1.435 |
| Ref2NM | 0.267 | | 0.453 | 0.028 | 0.095 | 0.60 | 3.535 | 1.319 |
| Ref2NM | 0.25 | | 0.431 | 0.022 | 0.01 | 0.60 | 3.539 | 1.238 |
| Ref2NM | 0.238 | | 0.443 | 0.022 | 0.098 | 0.60 | 3.55 | 1.349 |
| Ref2NM | 0.439 | | 0.418 | 0.054 | 0.0086 | 0.58 | 3.527 | 0.571 |
| Y14 | 0.40 | | 0.408 | 0.041 | 0.013 | 0.70 | 3.588 | 0.404 |
| Y14 | 0.216 | | 0.479 | 0.031 | 0.0014 | 0.51 | 3.692 | 1.697 |
| Y14 | 0.267 | | 0.453 | 0.028 | 0.095 | 0.60 | 3.624 | 1.148 |
| Y14 | 0.25 | | 0.431 | 0.022 | 0.01 | 0.60 | 3.618 | 1.276 |
| Y14 | 0.238 | | 0.443 | 0.022 | 0.098 | 0.60 | 3.636 | 1.064 |
| Y14 | 0.439 | | 0.418 | 0.054 | 0.0086 | 0.58 | 3.6 | 0.318 |

**Table S6. Heterotypic protein phase separation systems.**

| Protein | $N_{chains}$ | Conc [µM] | Box [nm] | LLPS in sim | Length [µs] | LLPS Reference |
|---|---|---|---|---|---|---|
| FUS LCD/ $(RGRGG)_5$ | 266/266 | 201/201 | 130 | No | 10 | Kaur *et al.*[19] |
| FUS LCD/ $(RGRGG)_5$ | 266/532 | 201/402 | 130 | No | 10 | Kaur *et al.*[19] |
| FUS LCD/ $(RGRGG)_5$ | 266/1330 | 201/1005 | 130 | Yes | 10 | Kaur *et al.*[19] |
| FUS LCD/ $(RGRGG)_5$ | 266/2660 | 201/2011 | 130 | Yes | 10 | Kaur *et al.*[19] |

**Table S7. Protein – RNA phase separation systems.**

| Protein | $N_{chains}$ | Conc [µM] | Box [nm] | LLPS in sim | Length [µs] | LLPS Reference |
|---|---|---|---|---|---|---|
| polyAde-21 / $(RRLR)_6$-SSSGSS | 126/147 | 210/240 | 100 | Yes | 10 | Bai *et al.*[23] |
| polyUra-40 / FUS $LCD_{RGG3}$ | 197/696 | 330/1160 | 100 | Yes | 4 | Kaur *et al.*[19] |
| polyUra-10 / polyArg-50 | 360/72 | 600/120 | 100 | Yes | 10 | Fisher & Elbaum-Garfinkle *et al.*[24] |
| polyAde-500 / $(RGRGG)_5$ | 30/1786 | 6/357 | 200 | No | 10 | Alshareedah *et al.*[25] |

**Table S8. Linear fit parameters for relating interaction energy to log($c_{sat}$) according to Eq. 9**

| Protein | a | b |
|---|---|---|
| hTau40-k18 | 8.88 | -11.64 |
| A1 LCD | 11.51 | -16.34 |
| LAF1 | 12.44 | -18.53 |
| RLP | 14.86 | -20.82 |
| FUS LCD | 13.19 | -21.61 |
| αSyn | 10.32 | -16.21 |
| hnRNPA1 | 20.96 | -75.51 |
| hSUMO_hnRNPA1 | 29.82 | -106.30 |
| MAGOH | 8.37 | -46.10 |
| Ref2NM | 33.00 | -115.86 |
| Y14 | 12.38 | -43.99 |

**Table S9. IDP $R_g$ test set**

| Protein | N | Rg [nm] | Reference |
|---|---|---|---|
| angiotensin | 8 | 0.79 | Ohnishi *et al.*[26] |
| ak16 | 16 | 0.98 | Kohn *et al.*[27] |
| Hist5 | 24 | 1.38 | Cragnell *et al.*[28] |
| CspTm | 67 | 1.47 | Müller-Späth *et al.*[29] |
| CTD2 | 83 | 2.61 | Gibbs *et al.*[30] |
| erm | 122 | 3.96 | Lens *et al.*[31] |
| A1 | 137 | 2.76 | Bremer *et al.*[16] |
| sNase | 141 | 2.12 | Flanagan *et al.*[32] |
| fhua | 142 | 3.34 | Riback *et al.*[33] |
| hTau-k25 | 185 | 4.1 | Mylonas *et al.*[34] |
| CAHSD | 229 | 4.8 | Hesgrove *et al.*[35] |
| hTau-k27 | 231 | 3.7 | Mylonas *et al.*[34] |
| PNt | 334 | 5.1 | Bowman *et al.*[36] |
| hTau-k25 | 450 | 4.1 | Mylonas *et al.*[34] |

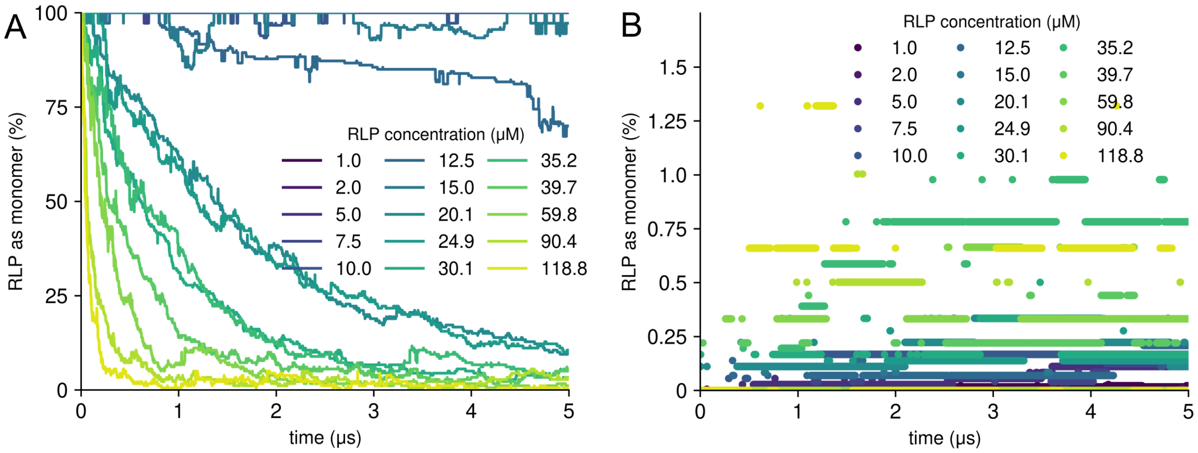


**Figure S1. Time evolution of RLP monomer fraction across various concentrations.** (A) and (B) show the percentage of RLP remaining as monomers over time at concentrations ranging from 1.0 μM to 118.8 μM starting from a randomly distributed (A) or condensate (B) state. Higher concentrations lead to faster depletion of monomers, indicating accelerated condensation.

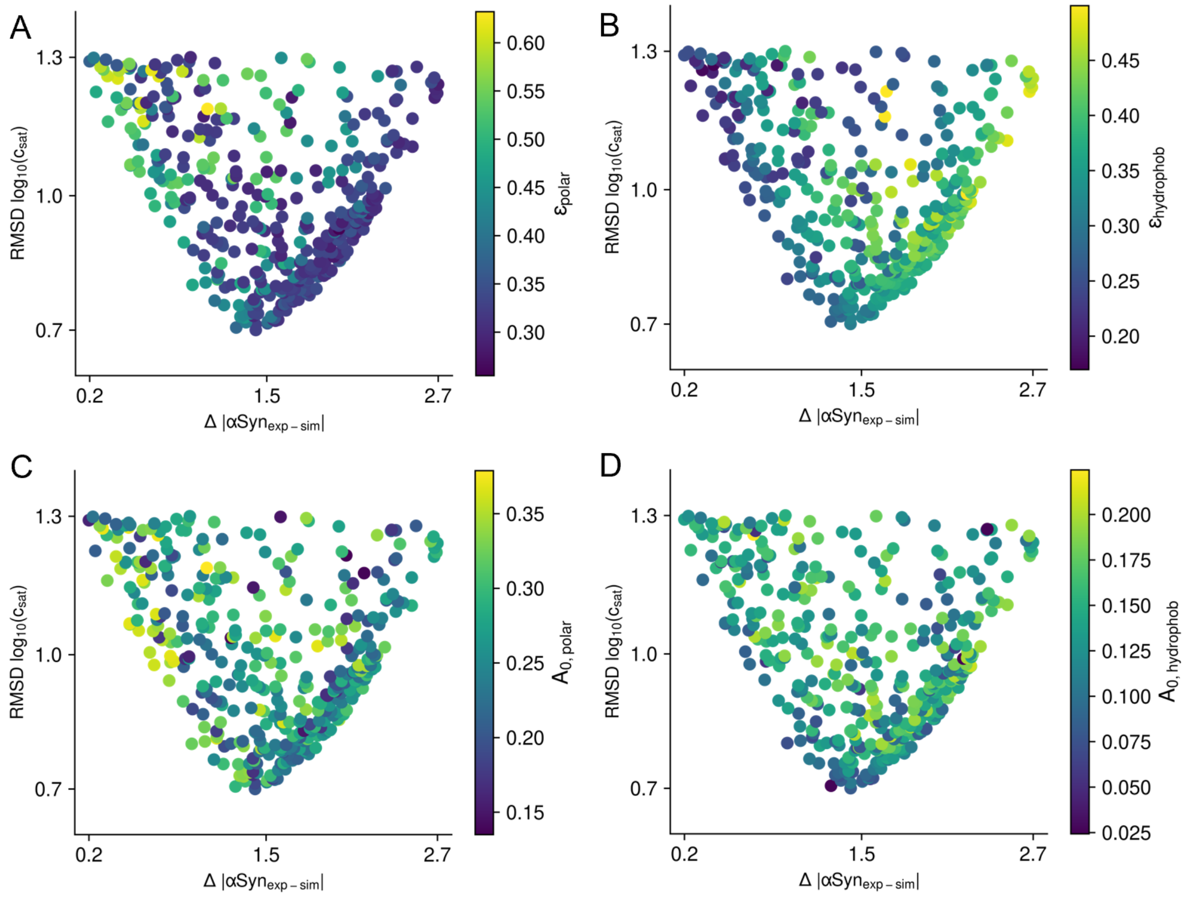


**Figure S2. Exploration of parameter space for COCOMO2.** (A-D) Comparison of RMSD deviation log10($c_{sat}$) for IDPs and the log10($c_{sat}$) deviation of αSyn from experimental values across 400 top-performing parameter sets. The points are colored based on the values of $\varepsilon_{polar}$ (A), $\varepsilon_{hydrophobic}$ (B), $A_{0,polar}$ (C) and $A_{0,hydrophobic}$ (D).

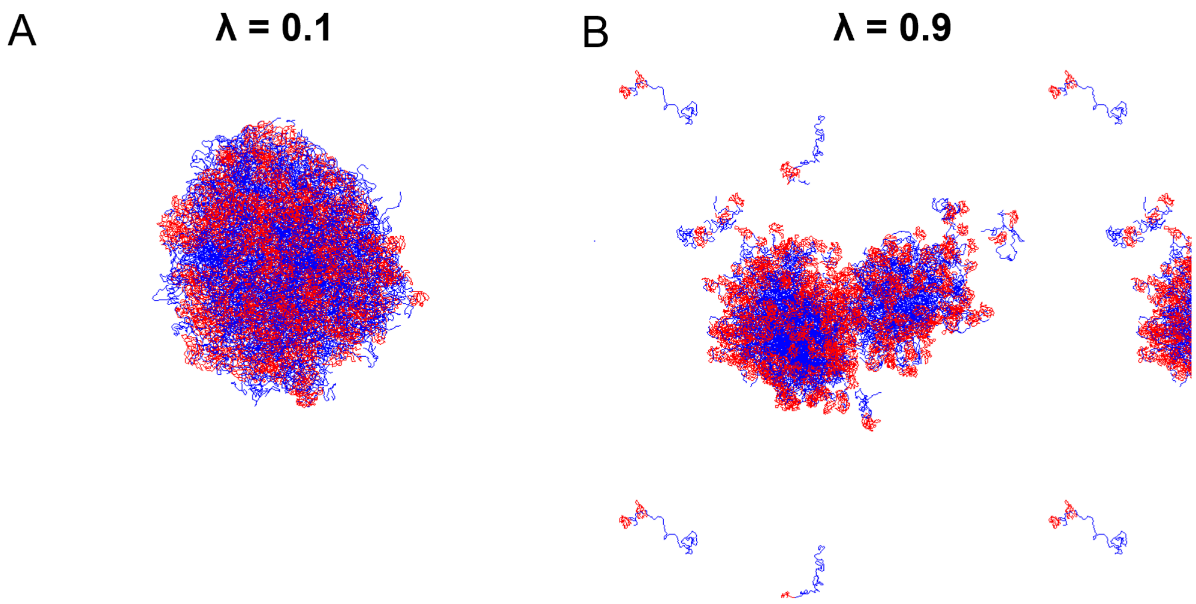


**Figure S3. Morphological change of hnRNPA1 condensates for different values of λ.** (A) At λ = 0.1, the folded domains are incorporated in the condensate, resulting in a dense structure. (B) At λ = 0.9, the folded domains become more flexible and are found at the surface of the condensate.

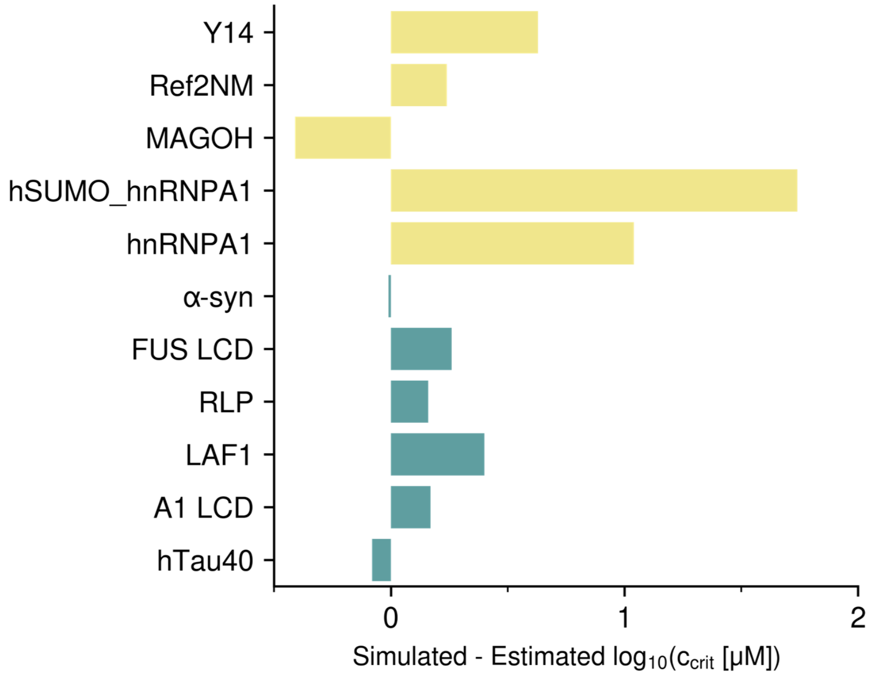


**Figure S4. Comparison of the difference between simulated and estimated $\log_{10}(c_{crit})$ values for IDPs and multi-domain proteins.** Proteins with smaller deviations are mostly IDPs (shown in blue), while larger deviations, particularly exceeding 1 log unit, are primarily observed for multi-domain proteins (in yellow).

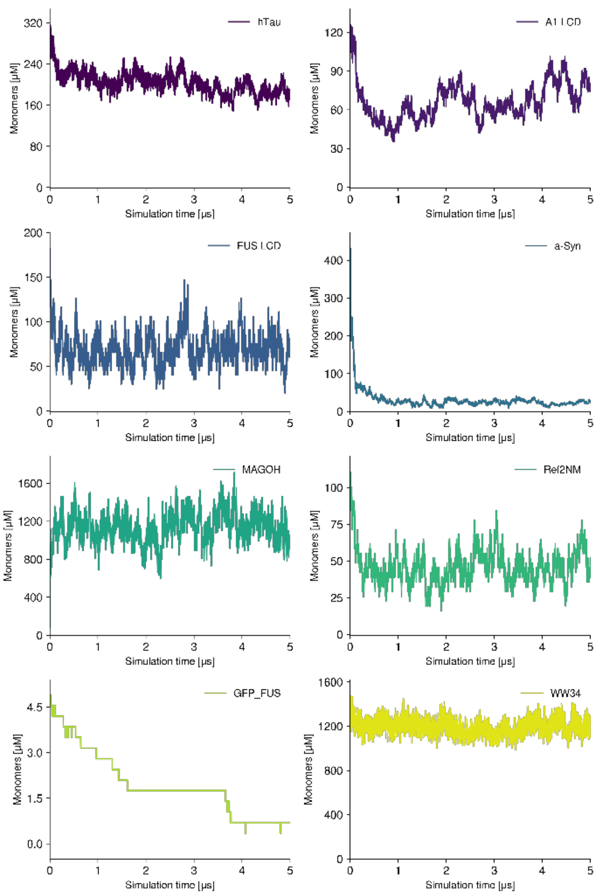


**Figure S5. Time evolution of monomer concentration across training and test systems for IDPs and multi-domain proteins.**

(25) Alshareedah, I.; Kaur, T.; Ngo, J.; Seppala, H.; Kounatse, L.-A. D.; Wang, W.; Moosa, M. M.; Banerjee, P. R. Interplay between Short-Range Attraction and Long-Range Repulsion Controls

Reentrant Liquid Condensation of Ribonucleoprotein–RNA Complexes. *J Am Chem Soc* **2019**, *141* (37), 14593–14602. https://doi.org/10.1021/jacs.9b03689.
